# Long-term Dietary Fat Intervention Affects Retinal Health in APOE Mice

**DOI:** 10.1101/2025.07.30.667671

**Authors:** Surabhi D. Abhyankar, Qianyi Luo, Gabriella D. Hartman, Neha Mahajan, Timothy W. Corson, Adrian L. Oblak, Bruce T. Lamb, Ashay D. Bhatwadekar

## Abstract

The *APOE* genotype influences metabolic and neurodegenerative outcomes, with *APOE4* carriers at higher risk for Alzheimer’s disease (AD) and metabolic dysfunction. This study examines how long-term dietary interventions affect systemic metabolism, retinal structure/ function in *APOE3*-knock-in (KI, neutral for AD) and *APOE4*-KI mice. Humanized *APOE3* and *APOE4*-KI mice received either a control diet (CD) or a Western diet (WD) for 2, 6, or 12 months. Body weight, glucose metabolism, lipid profiles, retinal structure, function, vasculature, visual performance, and inflammatory markers were analyzed. WD induced early glucose intolerance in *APOE4* mice (2 months); *APOE3* mice showed impairment only after prolonged exposure (6-12 months). Notably, WD-fed *APOE3* mice exhibited more pronounced hyperlipidemia than *APOE4* mice. *APOE4* CD mice displayed early retinal thinning (6 months), while *APOE4* WD mice initially exhibited retinal swelling, followed by degeneration (12 months). WD exacerbated retinal vascular dysfunction in *APOE4* mice, with increased tortuosity and reduced vascular area. Elevated *Il1b* expression in WD-fed *APOE4* mice confirmed inflammation-associated retinal dysfunction. *APOE4* mice showed heightened vulnerability to diet, with WD worsening metabolic, retinal, and vascular impairments. While CD improved glucose metabolism, it did not prevent retinal dysfunction. These findings underscore genotype-specific dietary strategies to mitigate *APOE4*-associated risks.

## Introduction

Alzheimer’s Disease (AD) is a major global health challenge, affecting over 55 million individuals worldwide, with the prevalence expected to rise as populations age.^1,2^ The most common form of AD, Late-Onset Alzheimer’s Disease (LOAD), typically manifests after the age of 65^3^, affecting nearly 30% of individuals over 85.^4,5^ The progression of AD is marked by cognitive decline, including memory loss, impaired learning, and executive dysfunction, ultimately hindering everyday activities.^6^

LOAD has a strong genetic component, with the apolipoprotein *E4* (*APOE4*) allele identified as the most significant genetic risk factor for disease onset and progression.^7^ The human *APOE* gene has three major alleles (*APOE2*, *APOE3*, and *APOE4*), with *APOE4* significantly increasing the risk of AD, while *APOE2* is associated with a protective effect, and *APOE3* is associated with a neutral effect.^8,9^ The presence of one or two copies of *APOE4* substantially accelerates the age of onset and exacerbates the severity of the disease.^10,11^ Studies have shown that *APOE4* not only contributes to cognitive decline but also influences brain metabolism and neuroinflammation, amplifying the pathological features of AD.^12–14^

Type 2 Diabetes Mellitus (T2D) and obesity are two prevalent metabolic disorders with significant overlap in pathology with AD, particularly in individuals carrying the *APOE4* allele. T2DM, which affects over 700 million people globally,^15^ is characterized by insulin resistance, impaired glucose regulation, and progressive organ dysfunction, including cognitive impairments.^16^ T2DM increases the risk of AD by 50-100%,^17–19^ with individuals suffering from both conditions exhibiting overlapping pathological features such as insulin resistance, oxidative stress, neuroinflammation,^19,20^ and amyloid plaque accumulation.^21,22^ Notably, T2D has been referred to as "type 3 diabetes" ^19,23,24^ due to the shared mechanisms between impaired glucose metabolism in the brain and the pathology of AD.^25^ The *APOE4* allele has been shown to exacerbate these metabolic dysfunctions, particularly in T2D, with studies demonstrating worsened neurodegeneration and cognitive decline in *APOE4* carriers who are diabetic.^26,27^

Obesity, a key risk factor for both T2D and AD,^28^ further amplifies these metabolic and cognitive disturbances, especially in *APOE4* carriers.^29^ Obese individuals with *APOE4* are at an even greater risk for developing T2DM and AD,^30,31^ with worsened insulin resistance, higher glucose levels, and cognitive decline.^32,33^ High-fat diets (HFDs), which mimic Western-style diets, are commonly used to induce obesity and glucose intolerance in experimental models, reflecting the pathophysiological features observed in individuals with T2DM and obesity.^34,35^ These diets serve as an effective tool for studying how metabolic disturbances, such as insulin resistance and obesity, contribute to retinal and brain dysfunction.

Given the significant overlap between T2D, obesity, and AD, the present study aims to explore the interplay between the *APOE4* allele, metabolic dysfunction, and retinal health. Using humanized *APOE3*-KI and *APOE4*-KI mouse models developed by the Model Organism Development and Evaluation for Late-Onset Alzheimer’s Disease (MODEL-AD) consortium, this study examines how different dietary interventions (particularly a Western style high-fat diet, referred to as WD) influence systemic glucose metabolism and retinal alterations, with a focus on the unique susceptibility associated with the *APOE4* allele. A recent study from our group demonstrated that middle-aged *APOE4*-KI mice exhibit notable retinal dysfunction, including structural, functional, and vascular deficits, which mirror the brain pathology observed in AD.^36^ The retina shares common features with the central nervous system,^37,38^ such as neural connectivity and vascular structure, making it an ideal model for studying the effects of metabolic dysfunction on brain health.^39,40^

This research investigates explicitly how a Western-style, high-fat diet, commonly associated with obesity and T2DM, exacerbates retinal changes in the presence of *APOE4*. We aim to determine whether the *APOE4* allele induces greater susceptibility to diet-induced retinal degeneration, even in the absence of overt diabetes, thereby offering a window into early pathophysiological changes. Here, we longitudinally assessed how the *APOE* genotype modulates systemic glucose metabolism, lipid profiles, and retinal structure-function in response to a Western-style diet, using humanized knock-in mice. Given the well-established connection between metabolic dysfunction and neurodegenerative diseases, this study aims to elucidate how *APOE4* exacerbates retinal and systemic metabolic disturbances, providing new insights into the intersection of obesity, T2D, and AD.

## Results

### Western diet induces glucose intolerance in *APOE4* mice

The effects of dietary intervention on body weight (BW) and glucose metabolism were first evaluated. *APOE3* and *APOE4* mice fed a WD exhibited a significant increase in BW after six months of treatment, whereas two months of WD exposure was insufficient to induce notable weight gain (Figure 1A). Glucose tolerance was assessed at multiple time points (2, 6, and 12 months), revealing unexpected findings. Despite no significant weight gain, *APOE4* mice on WD displayed marked glucose intolerance as early as two months post-treatment, in contrast to *APOE3* WD and *APOE4* CD groups (Figure 1B, C). By six months (Figure 1B, D) and twelve months (Figure 1B, E), glucose tolerance in *APOE4* WD mice became comparable to that of *APOE3* WD mice, suggesting that prolonged WD exposure in *APOE3* mice leads to systemic glucose dysregulation similar to that observed in *APOE4* mice.

**Figure 1.**
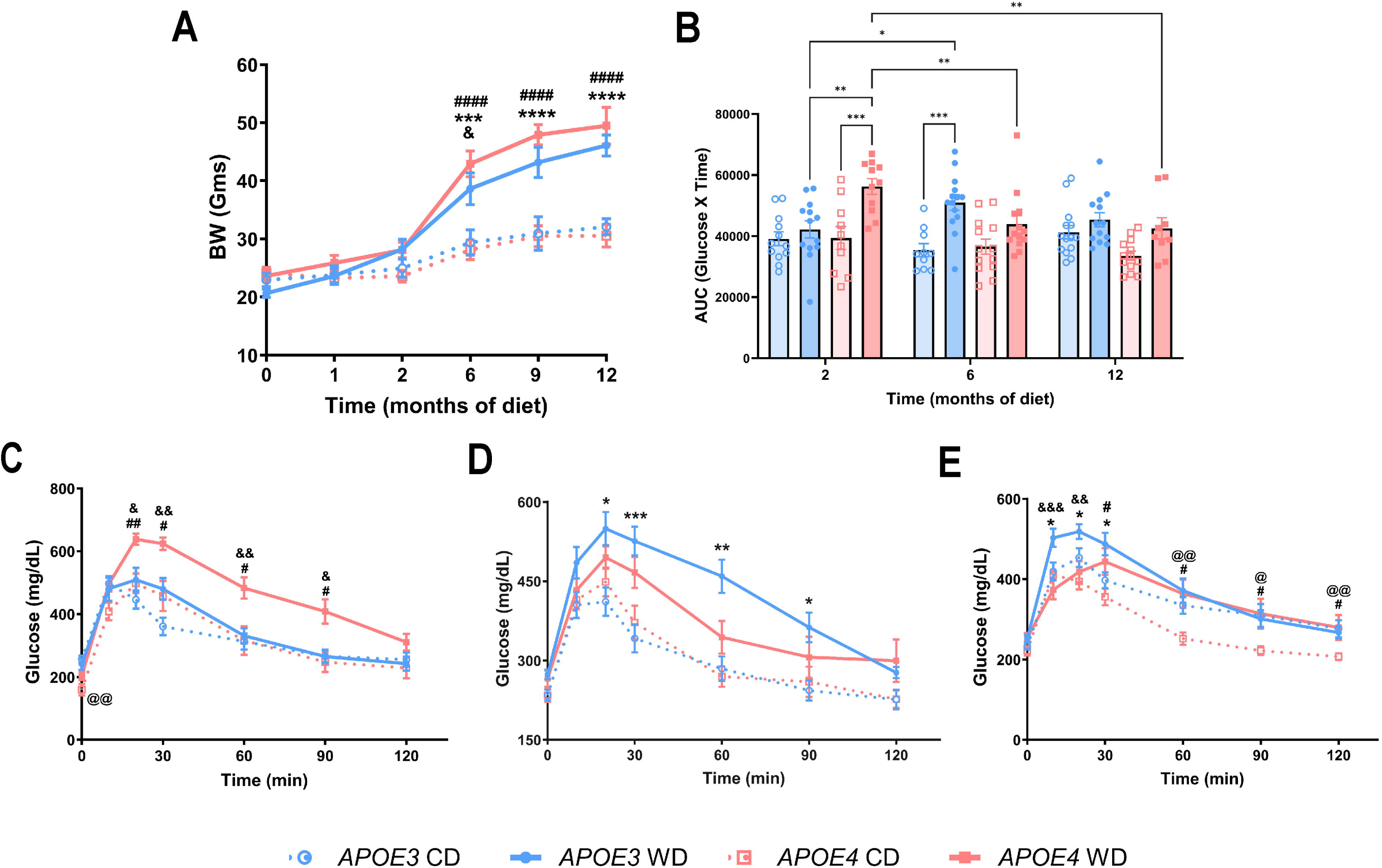
Effect of CD and WD on body weight and glucose tolerance. The impact of chronic dietary intervention on body weight (BW) and glucose metabolism was assessed over a 12-month period. (A) BW measurements show that WD significantly increased BW in both *APOE3* and *APOE4* mice compared to CD-fed controls, but substantial weight gain was only observed after 6 months of WD exposure. (B) Area under the curve (AUC) analysis of the intraperitoneal glucose tolerance test (IPGTT) at 2, 6, and 12 months shows that after prolonged WD exposure, *APOE3* mice develop comparable glucose intolerance to *APOE4* WD mice. (C) IPGTT at 2 months reveals that despite no significant weight gain, *APOE4* WD mice already exhibited marked glucose intolerance, while *APOE3* WD mice remained metabolically similar to their CD counterparts. (D, E) By 6 and 12 months, glucose intolerance worsened in *APOE3* WD mice, becoming similar to *APOE4* WD mice. Values are expressed as mean ± S.E.M. (n=13-14 mice/ group). Statistical analysis: Two-way ANOVA with Tukey’s test: **\***: *APOE3* CD vs *APOE3* WD, **@**: *APOE3* CD vs. *APOE4* CD, **&**: *APOE3* WD vs *APOE4* WD, **#**: *APOE4* CD vs *APOE4* WD). *p<0.05, **p<0.01, ***p<0.001, ****p<0.0001.

### Lipid panel reveals genotype-specific effects of WD on plasma lipids

To assess the systemic metabolic effects of chronic WD exposure, plasma lipid profiles were analyzed in *APOE3* and *APOE4* mice after 12 months on a CD or a WD. Overall, WD feeding resulted in increased plasma lipid levels in both genotypes, though the magnitude and pattern varied by genotype. Low-density lipoprotein (LDL, Supplemental Figure 1A) cholesterol levels were significantly elevated in WD-fed *APOE3* mice compared to their CD counterparts, while *APOE4* WD mice showed no significant change. Triglyceride (TG, Supplemental Figure 1B) levels trended upward in both genotypes with WD feeding but without reaching statistical significance. Similarly, total cholesterol (Supplemental Figure 1C) levels were elevated in both WD fed *APOE3* and *APOE4* mice, while WD-fed *APOE3* mice showed a statistically significant increase compared to CD-fed *APOE3* mice. Free fatty acid (FFA, Supplemental Figure 1D) levels displayed a divergent pattern. *APOE3* WD mice exhibited a reduction in FFA levels relative to *APOE3* CD controls, whereas *APOE4* WD mice showed significantly increased FFA levels compared to *APOE4* CD mice. Interestingly, *APOE3* CD mice also had elevated FFA levels compared to *APOE4* CD mice. High-density lipoprotein (HDL, Supplemental Figure 1E) cholesterol levels were significantly higher in *APOE3* WD mice compared to *APOE3* CD controls, with only modest, non-significant increases observed in *APOE4* WD mice. These results indicate that long-term WD exposure leads to systemic hyperlipidemia, particularly in *APOE3* mice, whereas *APOE4* mice demonstrate a more attenuated lipid response.

### Retinal structural alterations in *APOE4* mice following WD treatment

To investigate the impact of diet on retinal structure, OCT was employed to assess retinal thickness (Figure 2A). No significant differences were observed across groups after two months of dietary intervention. However, after six months, *APOE4* mice on CD exhibited a reduction in total retinal thickness (Figure 2B) and inner retinal thickness (Figure 2C). Interestingly, *APOE4* mice on WD demonstrated an increase in outer retinal thickness (Figure 2D), which may indicate retinal swelling induced by high-fat diet (HFD) consumption. By twelve months, outer retinal thickness in *APOE4* WD mice significantly decreased (Figure 2C), suggesting progressive retinal degeneration. Fundus imaging confirmed the absence of retinal lesions in all dietary groups (Figure 2A).

**Figure 2.**
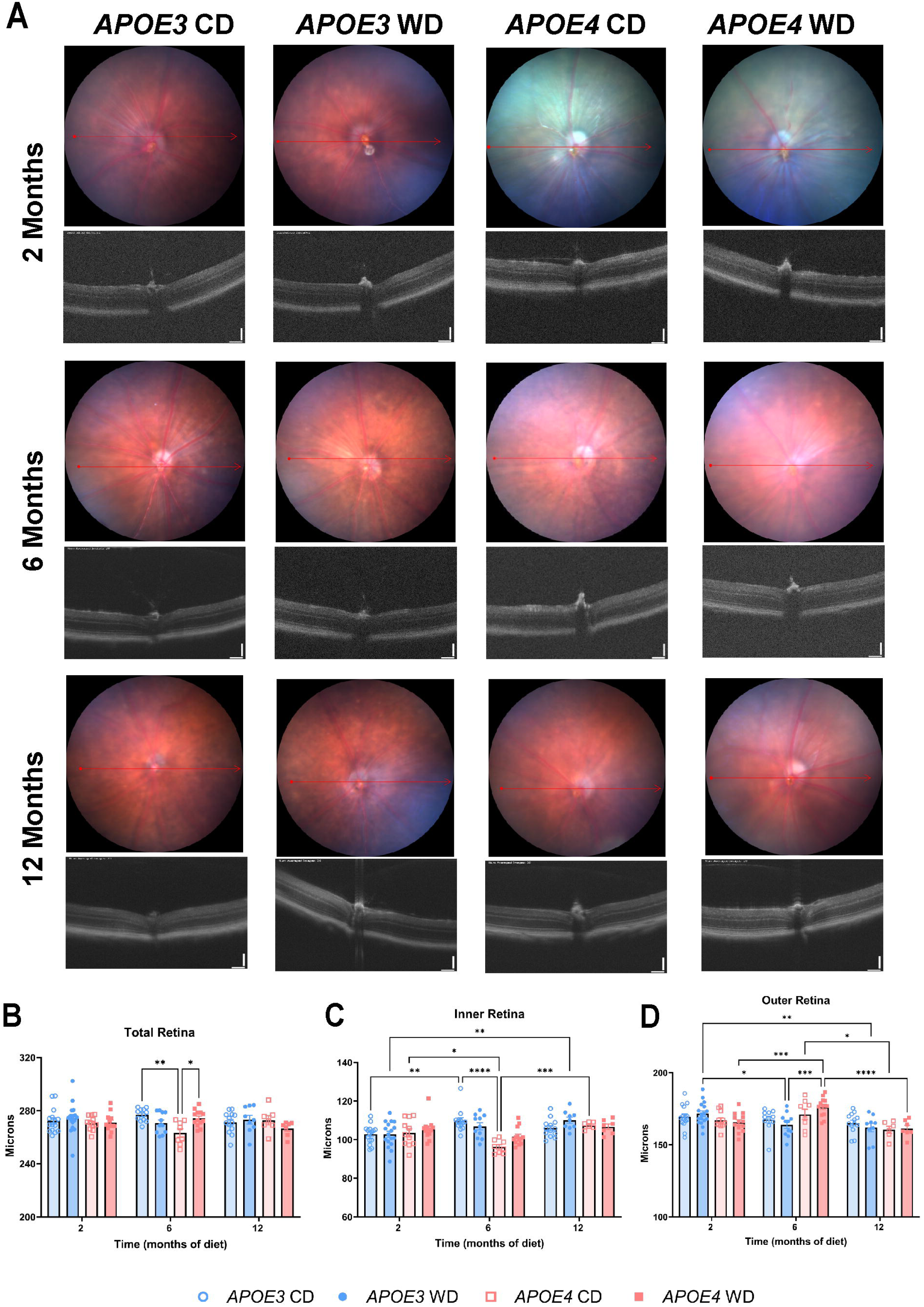
**Effect of CD and WD on retinal thickness**. Optical coherence tomography (OCT) was used to assess retinal thickness across diet and genotype groups. (A) Representative fundus and OCT images of *APOE3* and *APOE4* mice under CD and WD conditions at multiple time points (2, 6, and 12 months). No visible retinal lesions, hemorrhages, or gross abnormalities were detected across groups, suggesting that WD induced retinal dysfunction in *APOE4* mice occurs at a structural and functional level rather than through visible retinal damage. Scale bar: 100μm. n= 10-16 mice/ group. (B) Total retinal thickness was reduced in *APOE4* CD mice at 6 months, while *APOE4* WD mice initially exhibited increased thickness at 6 months, followed by significant thinning at 12 months, suggesting progressive degeneration. (C) Inner retinal thickness was reduced in *APOE4* CD mice at 6 months, indicating early structural deficits. (D) Outer retinal thickness increased in *APOE4* WD mice at 6 months, potentially due to retinal stress or swelling, but decreased by 12 months, suggesting photoreceptor loss. Values are expressed as mean ± S.E.M. (n= 10-16 mice/ group). Statistical analysis: Two-way ANOVA with Tukey’s test. *p<0.05, **p<0.01, ***p<0.001, ****p<0.0001.

### Western diet exacerbates retinal vascular impairments in *APOE4* mice

To determine whether WD compromises the integrity of the blood-retinal barrier in the presence of the *APOE4* allele, FA was conducted. Vascular abnormalities were evident as early as two months into the dietary intervention, with the *APOE4* genotype being more susceptible to these changes than *the APOE3 genotype*. By six months, WD-fed *APOE4* mice exhibited increased vascular tortuosity (Figure 3B), reduced vein width (Figure 3C), and decreased vascular area (Figure 3D). These deficits persisted at 12 months, with further increases in tortuosity, a reduction in vascular area, and an expansion of avascular regions (Figure 3E). Collectively, these findings indicate that WD exacerbates retinal vascular dysfunction in *APOE4* mice. Notably, vascular leakage was absent in all groups throughout the study duration (Figure 3A).

**Figure 3.**
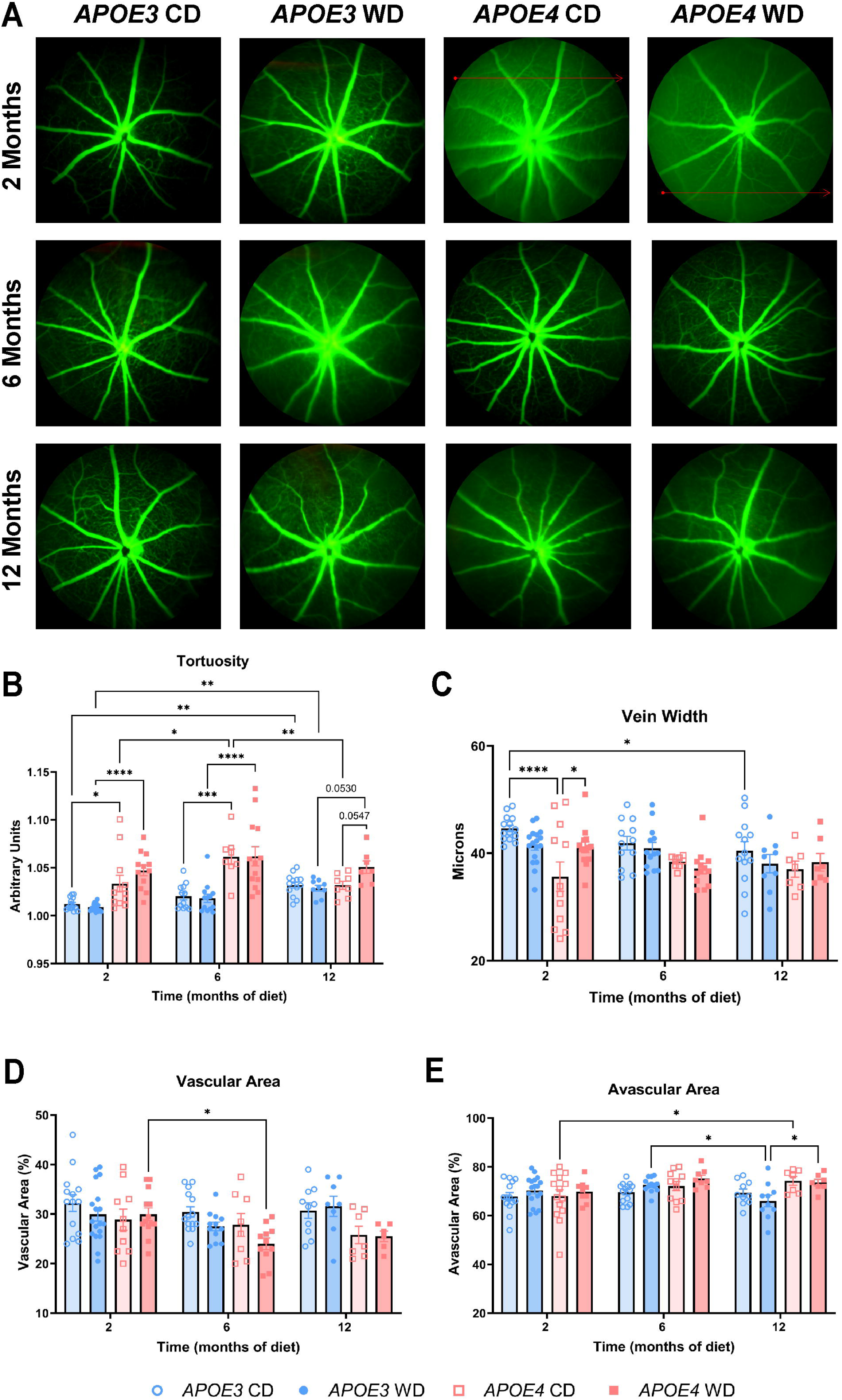
**WD exacerbates retinal vascular impairments in *APOE4* mice**. Fluorescein angiography (FA) was performed to assess vascular changes. (A) Vascular tortuosity increased in *APOE4* WD mice at 6 months, indicating progressive vessel abnormalities. (B) Vein width was reduced in *APOE4* CD mice at 2 months and further decreased in *APOE4* WD mice at 6 months. (C) The vascular area was significantly reduced in *APOE4* WD mice at 6 months, suggesting compromised retinal perfusion. (D) The avascular area increased in *APOE4* CD mice at 6 months, highlighting early microvascular deficits. Values are expressed as mean ± S.E.M. (n= 6-16 mice/ group). Statistical analysis: Two-way ANOVA with Tukey’s test. *p<0.05, **p<0.01, ****p<0.0001.

### Progressive decline in visual acuity and contrast sensitivity in WD-fed *APOE4* mice

To evaluate the functional consequences of dietary intervention on vision, OMR testing was performed. Following two months of WD exposure, *APOE4* mice exhibited significant reductions in both visual acuity (Figure 4A) and contrast sensitivity (Figure 4B) compared to *APOE3* WD and *APOE4* CD groups. These impairments progressed over the twelve months, with WD-fed *APOE4* mice displaying a continuous decline in both parameters. These results suggest that WD accelerates vision loss in *APOE4* mice, with the *APOE4* genotype conferring increased vulnerability to diet-induced visual impairments.

**Figure 4.**
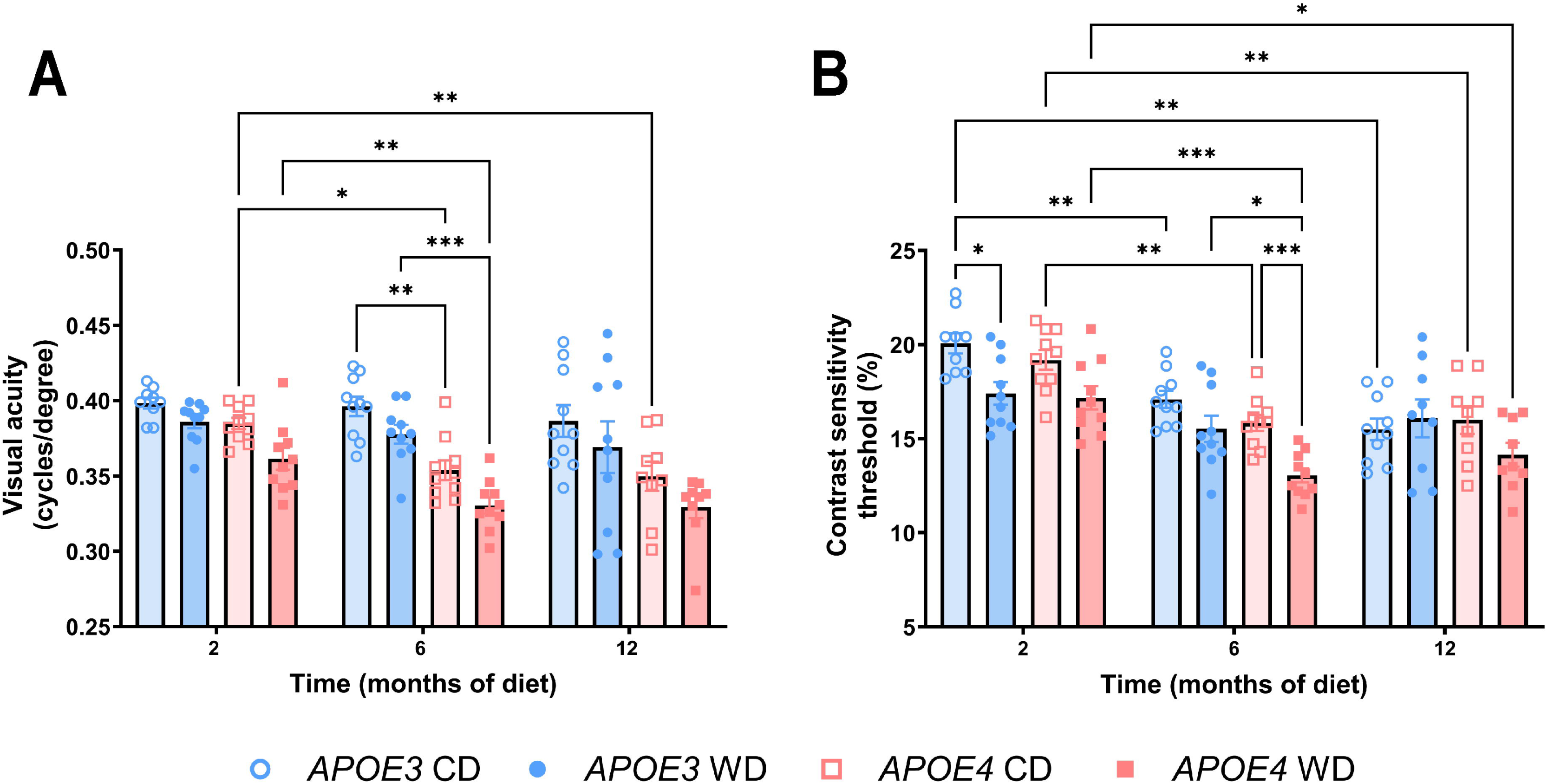
**Long-term WD results in lower visual acuity and contrast sensitivity in *APOE4* mice**. Optomotor response (OMR) testing was conducted to assess visual function. (A) Visual acuity measurements in APOE4 WD mice show a significant reduction as early as 2 months, which worsens progressively over the 12-month period, indicating a diet-induced decline in spatial vision. (B) Contrast sensitivity was also significantly reduced in WD fed *APOE4* mice, with impairments apparent at 2 months and further deteriorating over time. Values are expressed as mean ± S.E.M. (n= 9-15 mice/ group). Statistical analysis: Two-way ANOVA Tukey’s test. *p<0.05, **p<0.01, ***p<0.001.

### Retinal dysfunction in *APOE4* mice on CD and WD

Full-field ERG was utilized to assess retinal function under scotopic and photopic conditions. No significant differences in a or b-wave amplitudes or implicit times were observed among groups following two months of dietary treatment. However, by six and twelve months, both CD and WD-treated *APOE4* mice exhibited diminished scotopic a-wave (Figure 5A) and b-wave (Figure 5B) amplitudes. An increasing trend in a-wave implicit time was noted in these groups, although it did not reach statistical significance (Figure 5C). Additionally, WD-fed *APOE4* mice demonstrated a reduced b-wave implicit time under scotopic conditions at six months (Figure 5D). Under photopic conditions, *APOE4* CD mice showed significantly lower a-wave amplitudes (Figure 5E), while both CD and WD-fed *APOE4* mice exhibited significantly reduced b-wave amplitudes (Figure 5F). Furthermore, CD-fed *APOE4* mice demonstrated faster a-wave implicit times at six but not twelve months, despite having lower a-wave amplitudes. Meanwhile, WD-fed *APOE4* mice exhibited delayed a-wave implicit times (Figure 5G). Notably, both CD and WD-fed *APOE4* mice displayed faster b-wave implicit times (Figure 5H) despite reductions in amplitude. These results indicate that both diets impair the function of photoreceptors and bipolar cells in *APOE4* mice.

**Figure 5.**
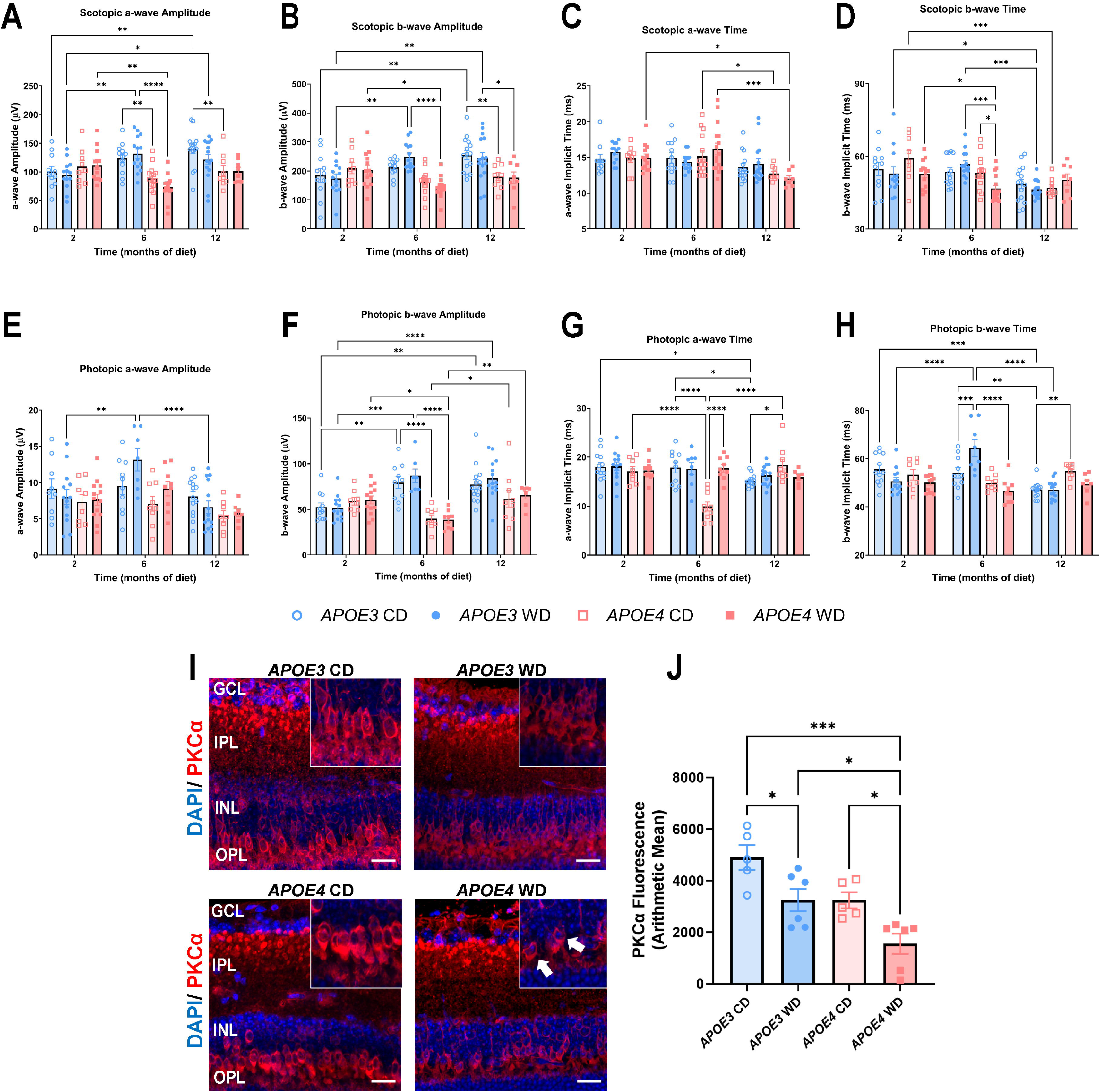
Impaired retinal function in *APOE4* mice on CD and WD. Electroretinography (ERG) was used to assess retinal function under scotopic and photopic conditions. (A, B) Scotopic a-wave and b-wave amplitudes were significantly reduced in *APOE4* CD and WD mice at 6 and 12 months, indicating progressive rod dysfunction. (C, D) Scotopic a-wave and b-wave implicit times were delayed in WD-fed *APOE4* mice, suggesting impaired signal transmission. (E, F) Photopic a-wave and b-wave amplitudes were reduced in both CD and WD fed *APOE4* mice, with greater deficits after 6 months, indicating cone dysfunction. (G, H) Photopic a-wave and b-wave implicit times were delayed in both CD and WD groups, further supporting retinal functional decline. To assess bipolar cell integrity, retinal sections were stained for PKCα, a marker of rod bipolar cells. (I) Representative images show bipolar cell stunting (white arrows) in *APOE4* WD mice, indicating structural abnormalities. Scale bar= 20 µm. (J) Quantification of PKCα fluorescence showed reduced bipolar cell labeling in WD fed *APOE4* mice, suggesting impaired bipolar cell integrity. Values are expressed as mean ± S.E.M. For ERG: n= 9-15 mice/ group, two-way ANOVA Tukey’s test. *p<0.05, **p<0.01, ***p<0.001, and ****p<0.0001. For PKCα: n= 3 mice/ group and 5-6 images/ group, one-way ANOVA with Tukey’s test. *p<0.05, ***p<0.001.

### Bipolar cell deficits in *APOE4* mice exposed to WD

Given the observed reduction in b-wave amplitudes in *APOE4* mice on CD and WD, bipolar cell integrity was further investigated via immunostaining for PKCα. Both CD and WD-treated *APOE4* mice exhibited a reduction in the density of bipolar cells (Figure 5I). Additionally, WD- fed *APOE4* mice demonstrated bipolar cell stunting (indicated by white arrows) along with a decrease in PKCα fluorescence intensity (Figure 5J). These findings suggest that while both dietary interventions disrupt bipolar cell function, WD further exacerbates cellular damage in *APOE4* mice.

### Western diet induces retinal inflammation in *APOE4* mice

*Il1b* is a key pro-inflammatory cytokine in both AD and T2D, where it contributes to neuronal injury, insulin resistance and vascular dysfunction.^41^ In our recent study we found elevated *Il1b* mRNA expression in *APOE4* retinas,^36^ therefore, we assessed effect of different dietary conditions on *Il1b* mRNA expression. qRT-PCR analysis revealed significantly elevated *Il1b* expression in WD-fed *APOE4* mice compared to both *APOE4*-CD and *APOE3*-WD groups (Figure 6A), indicating that chronic high-fat diet induces a genotype-specific pro-inflammatory response in the retina. Given the significant upregulation of *Il1b* in WD-fed *APOE4* mice, we next examined potential upstream and related pathways to better understand the inflammatory milieu. To investigate potential upstream drivers of *Il1b*, we examined *Nf-*κ*b*, a master regulator of inflammation activated by metabolic stress and associated with AD.^41^ In our study, *Nf-*κ*b* expression did not significantly differ across all groups (Figure 6B), indicating the *Il1b* elevation may occur through alternative or post-transcriptional mechanisms. *Vegfa* is involved in angiogenesis and vascular permeability; therefore, we assessed *Vegfa* expression to check whether WD-induced changes in retinal vasculature were accompanied by altered pro- angiogenic signaling. But no significant changes were observed in *Vegfa* expression across all groups (Figure 6C). Lastly, *Icam-1*, which is a marker for endothelial activation and leukocyte adhesion, which are elevated in metabolic disease and contribute to microvascular inflammation, did not show significant changes across all groups (Figure 6D). These findings put *Il1b* as the most sensitive marker for WD-induced retinal inflammation in *APOE4* mice.

**Figure 6.**
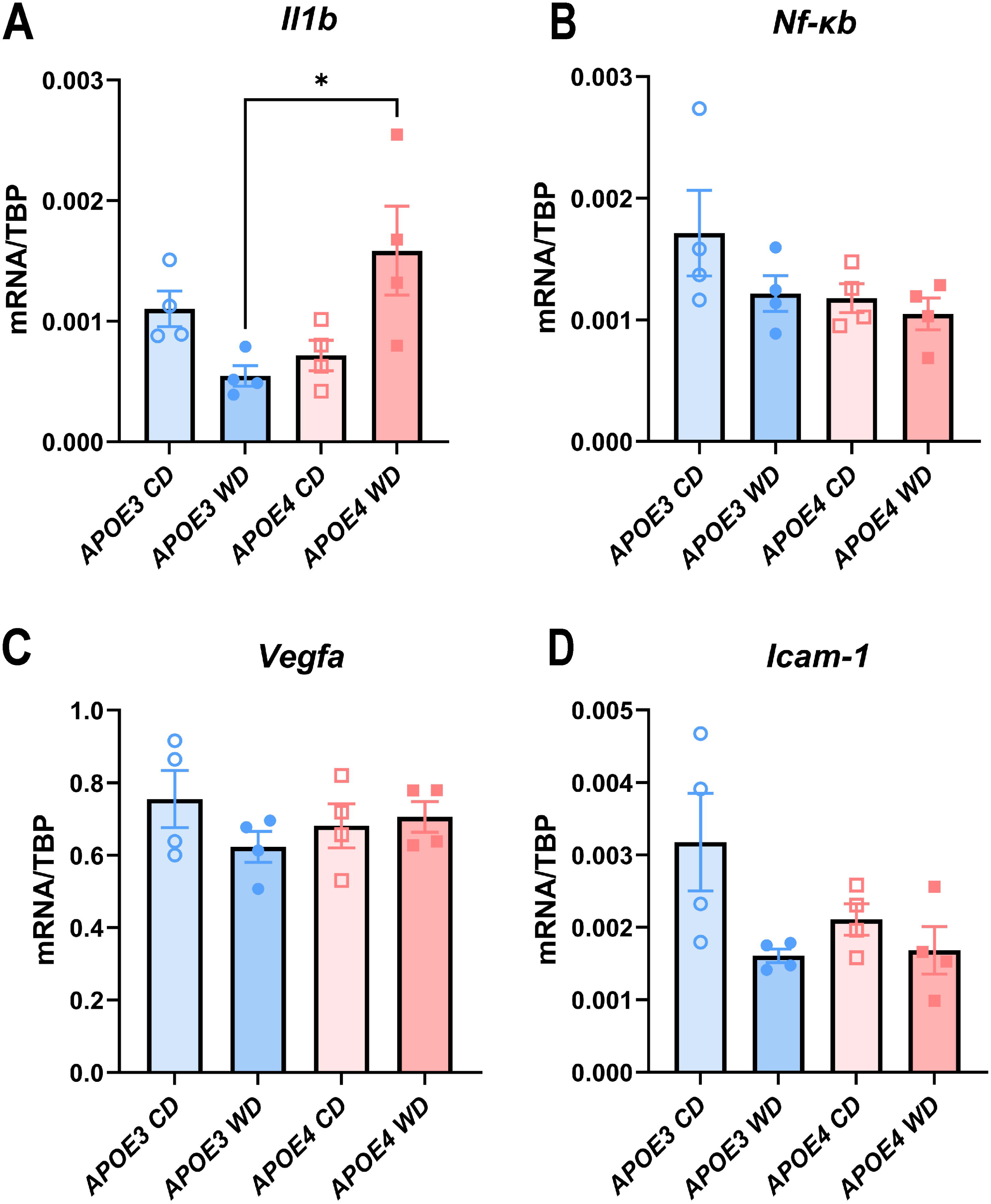
WD increases retinal *Il1b* expression and neuroinflammation in *APOE4* mice. (A) qRT-PCR from retinas showed significantly increased *Il1b* mRNA expression in WD fed *APOE4* mice, suggesting heightened neuroinflammatory responses. qRT-PCR data from additional inflammatory and vascular markers (B) *Nf-*κ*b,* (C) *Vegfa,* and (D) *Icam-1,* did not show significant changes across groups. Values are expressed as mean ± S.E.M. (n= 3-4 mice/ group). Statistical analysis: One-way ANOVA with Tukey’s test. *p<0.05.

## Discussion

This study investigates how the *APOE* genotype (*APOE3* and *APOE4*) influences systemic glucose metabolism, retinal integrity, vascular function, and visual performance in response to long-term dietary interventions. Using humanized *APOE*-KI mice, the findings highlight the heightened vulnerability of mice with the *APOE4* allele to dietary challenges, particularly high-fat diets (HFDs), which are associated with neurodegeneration and metabolic dysregulation.

The IPGTT results reveal distinct genotype-dependent metabolic responses to diet. While *APOE3* and *APOE4* mice had similar BW, WD-fed *APOE4* mice exhibited glucose intolerance after 2 months, aligning with previous studies.^42^ However, after 6 months, *APOE3* WD-fed mice showed greater glucose impairment, and by 12 months, *APOE4* mice demonstrated better glucose tolerance on a CD than on WD. This challenges the conventional link between *APOE4* and systemic glucose dysregulation^43–45^ and suggests a possible protective role of CD in *APOE4* mice. The observed metabolic resilience of *APOE4* compared to *APOE3* under WD conditions supports prior findings that *APOE3* mice develop systemic metabolic dysfunction, whereas *APOE4* mice primarily exhibit cerebral glucose deficits.^46^ Given that cerebral metabolic disturbances are implicated in AD progression,^47^ these genotype-specific metabolic responses warrant further exploration.

Diet composition appears to critically influence weight gain patterns in *APOE3* mice. Prior studies show that *APOE3*, rather than *APOE4*, mice gain BW when exposed to HFDs.^33,48^ The discrepancy in weight gain across studies may stem from variations in fat composition; this study used a milk-based 40% kcal fat diet, whereas others employed 45% kcal fat^42^ or 60% kcal fat from lard.^33^ These findings underscore that dietary fat type and proportion may distinctly impact *APOE4*, paralleling observations in human *APOE4* carriers who exhibit lower body mass index yet heightened susceptibility to metabolic and cognitive impairments.^49^

The lipid panel data show a clear genotype-specific metabolic response to chronic WD feeding. Contrary to expectations based on *APOE4*’s known association with dyslipidemia and AD risk in humans^50^, WD-fed *APOE3* mice exhibited a more severe hyperlipidemic phenotype following WD exposure. Our findings of elevated cholesterol and TG levels in WD-fed *APOE3* mice are consistent with the study by Huebbe *et al*. (2024), ^51^, which demonstrated that *APOE3* promotes hepatic lipid accumulation, leading to more severe fatty liver disease compared to *APOE4*. This shared lipid phenotype reinforces the notion that *APOE3* may contribute to systemic lipid dysregulation under HFD conditions, in contrast to the relatively blunted lipid response seen in *APOE4* mice. These findings emphasize that systemic lipid elevations are not the sole determinant of disease risk, particularly in WD-fed *APOE4* mice, and highlight the importance of exploring genotype-dependent lipid trafficking and compartmentalization, including within the retina and central nervous system.

OCT analyses reveal that dietary fat influences retinal morphology. After 6 months, CD- fed *APOE4* mice exhibited retinal thinning, while WD-fed *APOE4* mice displayed increased outer retinal thickness, indicative of retinal swelling. This suggests that *APOE4* mice experience significant retinal structural changes in response to dietary variations. Vascular analysis further highlights the detrimental effects of WD in *APOE4* mice, with increased vascular tortuosity, narrowed veins, and a reduction in vascular area observed after 6 and 12 months. While no vascular leakage was detected, these vascular abnormalities suggest WD- induced vascular dysfunction, potentially exacerbating blood-retinal barrier disruption.^52^ The lack of leakage does not rule out subtle barrier compromise or altered endothelial signaling, and future studies might consider direct assessments of permeability or tight junction integrity.

Visual function assessments reinforce the heightened sensitivity of *APOE4* mice to WD. OMR testing revealed reduced visual acuity and contrast sensitivity in WD-fed *APOE4* mice as early as 2 months, persisting through 12 months. Systemic and retinal lipid imbalances are well-established contributors to retinal degeneration.^53,54^ In this study, a diet comprising 40% of calories from fat was found to be less detrimental to retinal health than the more pronounced damage observed with a diet providing 60% of calories from fat. To ensure caloric equivalence when lowering fat intake, the carbohydrate content in the CD was increased from 50gm per kg (43% of calories from carbohydrates) to 71gm per kg (73% of calories from carbohydrates), with an additional 645g/kg of corn starch incorporated to balance the caloric intake of the WD. However, this modification resulted in a carbohydrate-dense diet, which may introduce its own set of adverse effects. Prior research similarly suggests that excessive carbohydrate intake can lead to retinal dysfunction.^55,56^ These findings suggest that while prolonged WD consumption exacerbates photoreceptor and bipolar cell dysfunction, extreme reductions in dietary fat coupled with excessive carbohydrate intake may also negatively impact retinal integrity. Furthermore, WD-fed *APOE4* mice exhibited delayed b-wave implicit times, signifying progressive retinal dysfunction. Notably, bipolar cell stunting in WD-fed *APOE4* mice provides a mechanistic explanation for these functional deficits, indicating diet-induced neurodegeneration.

Neuroinflammatory markers further support the role of diet in exacerbating *APOE4*- associated retinal damage. Elevated *Il1b* mRNA expression in WD-fed *APOE4* mice confirms inflammation-driven retinal dysfunction, even though other canonical markers such as *Nf-*κ*b, Icam1*, and *Vegfa* did not show consistent changes in this context. This may reflect stage- specific or region-specific activation not captured at the mRNA level, or simply that *Il1b* is a more sensitive early marker in this model. These findings illustrate that while CD may confer metabolic benefits, it does not necessarily protect against retinal dysfunction, emphasizing the need for balanced dietary approaches in *APOE4* carriers.

The critical part of interpreting our findings involves distinguishing between different dietary components. Unlike grain-based regular chow diets (NCDs) used in prior studies,^57^ including our own recently published work, which showed more pronounced retinal deficits in *APOE4* mice,^36^, the CD and WD in the present study are purified and compositionally defined.^58^ This compositional precision eliminates confounders inherent in NCDs, such as variable plant-derived compounds, complex macronutrient mixes, and fiber variability, allowing a more controlled assessment of *APOE*-diet interactions. As detailed in our methods, the CD is high in carbohydrates (73% kcal, primarily corn starch and maltodextrin) and low in fat (10% kcal). In contrast, the WD is high in fat (40% kcal, predominantly milk fat) and contains significant sucrose (341 g/kg). This distinction is critical, as the quantity and quality of carbohydrates significantly influence metabolic and inflammatory outcomes (Ludwig et al., 2018). While our previous NCD study consistently highlighted APOE4 vulnerability, the current results demonstrate that *APOE4*-related retinal deficits are not uniformly worse, but rather dynamically shift with the specific dietary environment. this is not contradictory; rather, it highlights how significantly different base diets can influence phenotype expression. The observed milder phenotype in *APOE4*-CD mice compared to our prior NCD report, and the exacerbation of deficits by WD, strongly suggest that *APOE4*-driven disease risk is not fixed, but substantially modifiable by the precise composition and quality of the dietary environment.

The findings from our study are supported by a full set of longitudinal data (mean ± SD) across metabolic, retinal, vascular, and inflammatory parameters detailed in table 1. These outcomes are also visually summarized in Figure 7, which summarizes the combined effects of *APOE* genotype and long-term diet on systemic and retinal health at 12 months. These findings hold several key implications: 1. Diet as a Modifiable Factor: The differential effects of CD and WD in *APOE4* mice suggest that dietary interventions can either mitigate or amplify retinal and metabolic impairments in *APOE4* carriers. 2. Genotype-Specific Responses: The contrasting responses of *APOE3* and *APOE4* mice to dietary fat emphasize the importance of personalized nutrition based on genetic background. 3. Pathophysiological Mechanisms: The complex interplay between *APOE4*, systemic metabolism, and neuroinflammation underscores the need for further research into how *APOE4* amplifies diet-induced damage.

**Figure 7.**
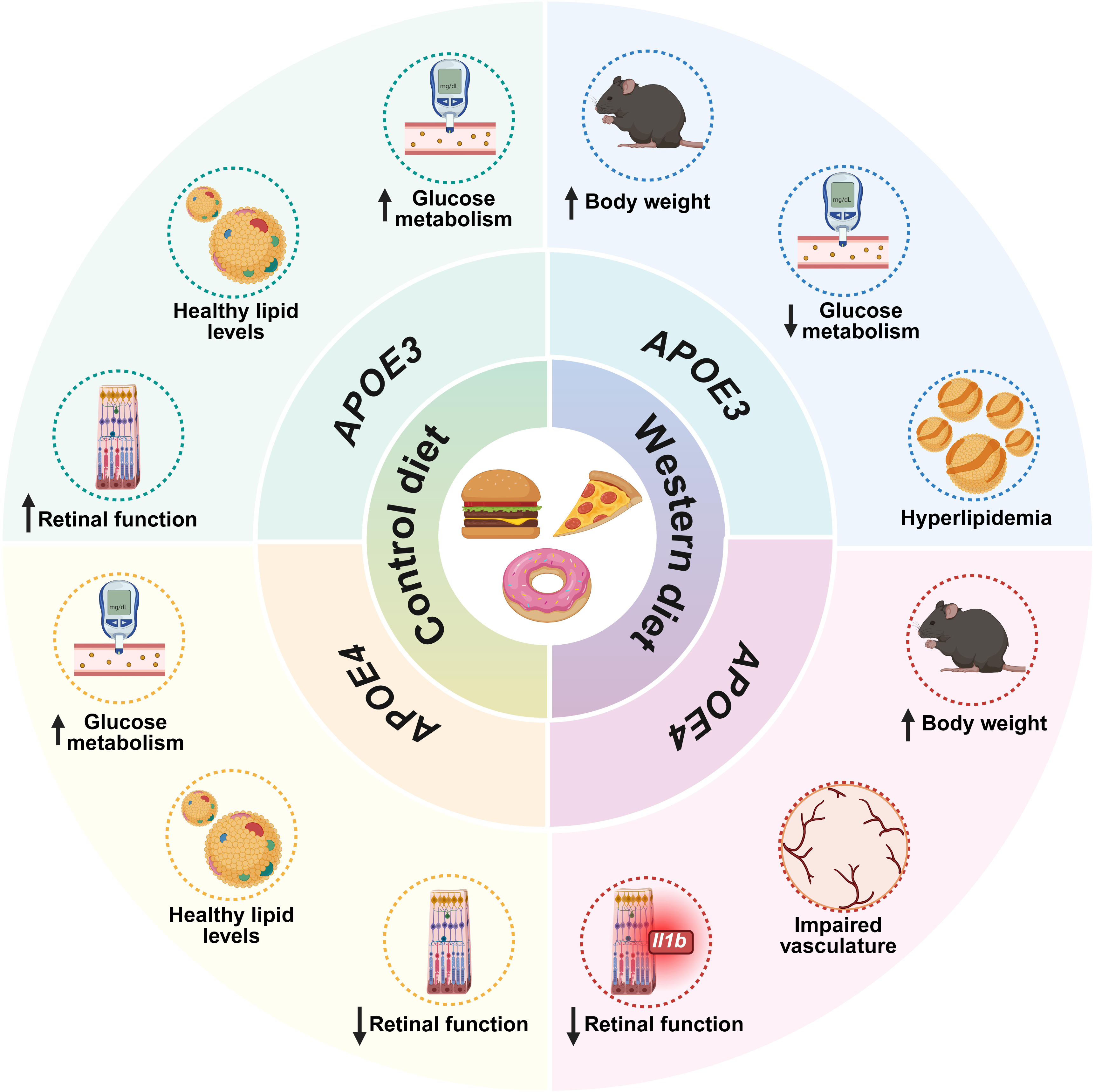
12-month impact of diet and *APOE* genotype on systemic and retinal health This schematic illustration summarizes the effects of long-term dietary intervention on metabolic and retinal health in *APOE*-KI mice at 12 months. The circular layout contrasts control diet (CD, left panels) and a Western diet (WD, right panels) outcomes across *APOE3* (top half) and *APOE4* (bottom half) genotypes. *APOE3*-CD mice exhibited improved glucose metabolism and retinal function with healthy lipid levels. In contrast, *APOE3*-WD mice showed increased body weight, hyperlipidemia, and reduced glucose metabolism. *APOE4*-CD mice maintained metabolic health but showed reduced retinal function. *APOE4*-WD mice displayed compounded dysfunction, including increased body weight, impaired retinal vasculature, and reduced retinal function with inflammatory signatures (increased *Il1b*), and signs of neurovascular compromise. Icons depict key phenotypes such as retinal function, vascular integrity, glucose metabolism, lipid status, and inflammatory signals. The figure highlights how WD induces widespread dysfunction in both genotypes, with APOE4 conferring additional susceptibility to retinal and vascular damage.

**Table 1:** Time-course effects of CD and WD on metabolic, retinal, and neuroinflammatory parameters in *APOE3* and *APOE4* mice.

| Parameters | APOE3 CD |  |  |  |  |  | APOE3 WD |  |  |  |  |  | APOE4 CD |  |  |  |  |  | APOE4 WD |  |  |  |  |  |
| --- | --- | --- | --- | --- | --- | --- | --- | --- | --- | --- | --- | --- | --- | --- | --- | --- | --- | --- | --- | --- | --- | --- | --- | --- |
|  | 2 months |  | 6 months |  | 12 months |  | 2 months |  | 6 months |  | 12 months |  | 2 months |  | 6 months |  | 12 months |  | 2 months |  | 6 months |  | 12 months |  |
|  | Mea<br>n | SD | Mea<br>n | SD | Mea<br>n | SD | Mea<br>n | SD | Mea<br>n | SD | Mea<br>n | SD | Mea<br>n | SD | Mea<br>n | SD | Mea<br>n | SD | Mea<br>n | SD | Mea<br>n | SD | Mea<br>n | SD |
| BW | 24.9<br>7 | 5.6<br>3 | 29.3<br>9 | 6.5<br>9 | 32.0<br>9 | 5.2<br>0 | 28.2<br>4 | 7.03 | 38.6<br>5 | 8.7<br>0 | 46.0<br>8 | 6.5<br>7 | 23.6<br>1 | 3.72 | 28.1<br>4 | 6.1<br>2 | 30.5<br>7 | 7.0<br>3 | 28.1<br>0 | 5.0<br>0 | 42.9<br>1 | 8.34 | 49.4<br>7 | 9.52 |
| GTT AUC | 391<br>24.5<br>8 | 786<br>6.7<br>5 | 354<br>11.5<br>0 | 676<br>4.8<br>0 | 413<br>07.5<br>0 | 825<br>9.3<br>7 | 422<br>31.9<br>2 | 101<br>89.0<br>6 | 510<br>49.2<br>9 | 947<br>4.1<br>1 | 453<br>90.7<br>7 | 824<br>0.5<br>6 | 393<br>61.0<br>0 | 119<br>17.2<br>3 | 365<br>40.7<br>7 | 891<br>0.4<br>3 | 335<br>96.9<br>2 | 531<br>3.3<br>6 | 562<br>50.0<br>0 | 851<br>4.1<br>6 | 439<br>43.8<br>5 | 103<br>86.3<br>1 | 425<br>78.8<br>9 | 104<br>11.0<br>9 |
| LDL | - | - | - | - | 8.46 | 2.1<br>2 | - | - | - | - | 48.9<br>0 | 26.<br>48 | - | - | - | - | 10.7<br>4 | 5.7<br>2 | - | - | - | - | 24.7<br>5 | 13.8<br>1 |
| TG | - | - | - | - | 107.<br>50 | 13.<br>80 | - | - | - | - | 175.<br>60 | 67.<br>16 | - | - | - | - | 78.1<br>4 | 22.<br>18 | - | - | - | - | 139.<br>93 | 86.1<br>4 |
| Cholesterol | - | - | - | - | 88.8<br>7 | 31.<br>86 | - | - | - | - | 246.<br>02 | 93.<br>67 | - | - | - | - | 73.3<br>8 | 9.4<br>4 | - | - | - | - | 194.<br>41 | 93.8<br>4 |
| FFA | - | - | - | - | 0.88<br>5 | 0.1<br>46 | - | - | - | - | 0.63<br>6 | 0.1<br>08 | - | - | - | - | 0.31<br>2 | 0.1<br>16 | - | - | - | - | 0.72<br>0 | 0.27<br>0 |
| HDL | - | - | - | - | 69.7<br>4 | 32.<br>84 | - | - | - | - | 152.<br>19 | 62.<br>33 | - | - | - | - | 50.5<br>0 | 17.<br>00 | - | - | - | - | 130.<br>42 | 53.4<br>0 |
| Vein width | 44.6<br>2 | 2.3<br>2 | 41.8<br>9 | 4.4<br>0 | 40.4<br>6 | 6.1<br>1 | 41.3<br>9 | 3.38 | 40.9<br>3 | 4.0<br>4 | 38.0<br>6 | 5.1<br>0 | 35.6<br>3 | 9.45 | 38.4<br>1 | 1.1<br>1 | 37.0<br>4 | 3.9<br>1 | 41.1<br>1 | 3.7<br>1 | 37.1<br>3 | 3.52 | 38.3<br>3 | 4.34 |
| Tortuosity | 1.01<br>22 | 0.0<br>065 | 1.02<br>03 | 0.0<br>131 | 1.03<br>21 | 0.0<br>115 | 1.00<br>88 | 0.00<br>42 | 1.01<br>82 | 0.0<br>157 | 1.02<br>88 | 0.0<br>090 | 1.03<br>35 | 0.02<br>93 | 1.06<br>14 | 0.0<br>232 | 1.03<br>18 | 0.0<br>119 | 1.04<br>72 | 0.0<br>194 | 1.06<br>19 | 0.03<br>66 | 1.05<br>07 | 0.01<br>74 |
| Vascular area | 32.2<br>1 | 6.3<br>5 | 30.4<br>3 | 3.9<br>7 | 30.7<br>0 | 4.9<br>7 | 29.9<br>4 | 5.67 | 27.5<br>0 | 2.9<br>9 | 31.5<br>6 | 5.7<br>4 | 28.8<br>5 | 6.74 | 27.8<br>1 | 6.5<br>5 | 25.7<br>9 | 4.6<br>7 | 29.9<br>6 | 4.5<br>8 | 24.0<br>0 | 3.84 | 25.5<br>0 | 2.59 |
| Avascular area | 67.7<br>9 | 6.3<br>5 | 69.5<br>7 | 3.9<br>7 | 69.3<br>0 | 4.9<br>7 | 70.2<br>2 | 5.79 | 72.5<br>0 | 2.9<br>9 | 66.0<br>0 | 7.3<br>7 | 67.9<br>3 | 9.69 | 72.0<br>9 | 6.0<br>2 | 74.2<br>1 | 4.6<br>7 | 69.7<br>8 | 4.5<br>2 | 75.1<br>9 | 3.87 | 73.5<br>0 | 3.55 |
| Total retina | 272.<br>27 | 9.1<br>6 | 276.<br>74 | 4.8<br>2 | 271.<br>10 | 7.9<br>9 | 274.<br>31 | 11.3<br>4 | 270.<br>70 | 7.8<br>3 | 273.<br>17 | 9.9<br>1 | 270.<br>54 | 5.33 | 263.<br>22 | 8.7<br>9 | 272.<br>79 | 8.9<br>1 | 270.<br>96 | 6.8<br>3 | 274.<br>25 | 6.03 | 266.<br>57 | 5.54 |
| Inner retina | 102.<br>84 | 4.8<br>6 | 109.<br>71 | 6.0<br>5 | 105.<br>96 | 4.6<br>9 | 102.<br>74 | 6.52 | 106.<br>88 | 6.0<br>2 | 110.<br>08 | 4.8<br>3 | 103.<br>51 | 7.01 | 96.2<br>6 | 3.4<br>4 | 107.<br>36 | 1.8<br>2 | 105.<br>40 | 5.5<br>6 | 101.<br>58 | 4.52 | 106.<br>65 | 3.68 |
| Outer retina | 169.<br>43 | 7.9<br>2 | 167.<br>50 | 7.7<br>6 | 165.<br>15 | 6.9<br>9 | 171.<br>57 | 7.79 | 163.<br>83 | 8.0<br>1 | 161.<br>98 | 8.8<br>3 | 167.<br>02 | 6.08 | 171.<br>09 | 10.<br>08 | 160.<br>42 | 5.9<br>6 | 165.<br>57 | 6.6<br>0 | 176.<br>24 | 6.48 | 161.<br>21 | 5.93 |
| Visual acuity | 0.39<br>8 | 0.0<br>11 | 0.39<br>6 | 0.0<br>20 | 0.38<br>6 | 0.0<br>33 | 0.38<br>6 | 0.01<br>4 | 0.37<br>8 | 0.0<br>20 | 0.36<br>9 | 0.0<br>54 | 0.38<br>5 | 0.01<br>2 | 0.35<br>4 | 0.0<br>21 | 0.35<br>0 | 0.0<br>29 | 0.36<br>1 | 0.0<br>23 | 0.33<br>0 | 0.01<br>7 | 0.32<br>9 | 0.02<br>2 |
| Contrast sensitivity | 20.0<br>7 | 1.6<br>3 | 17.0<br>8 | 1.4<br>1 | 15.4<br>9 | 1.8<br>1 | 17.3<br>9 | 1.91 | 15.5<br>2 | 2.2<br>1 | 16.0<br>8 | 3.0<br>1 | 19.1<br>8 | 1.66 | 15.8<br>3 | 1.3<br>5 | 15.9<br>9 | 2.2<br>2 | 17.1<br>7 | 1.9<br>5 | 13.0<br>5 | 1.19 | 14.1<br>4 | 1.88 |
| Scotopic a-wave amplitude | 100.<br>63 | 29.<br>75 | 123.<br>70 | 26.<br>56 | 139.<br>57 | 31.<br>39 | 94.9<br>8 | 25.9<br>9 | 131.<br>77 | 30.<br>53 | 121.<br>27 | 34.<br>83 | 109.<br>17 | 30.7<br>6 | 88.4<br>3 | 23.<br>25 | 101.<br>48 | 29.<br>58 | 111.<br>39 | 28.<br>00 | 73.2<br>4 | 20.7<br>0 | 101.<br>30 | 19.2<br>9 |
| Scotopic a-wave time | 14.7<br>5 | 2.1<br>2 | 14.9<br>2 | 2.5<br>1 | 13.6<br>2 | 1.9<br>6 | 15.7<br>7 | 1.37 | 14.3<br>9 | 1.2<br>0 | 14.1<br>0 | 2.7<br>7 | 14.8<br>6 | 1.81 | 15.2<br>0 | 2.8<br>2 | 12.7<br>5 | 0.9<br>7 | 14.9<br>6 | 1.8<br>8 | 16.1<br>9 | 3.28 | 12.0<br>4 | 1.11 |
| Scotopic b-wave amplitude | 185.<br>53 | 79.<br>36 | 212.<br>93 | 29.<br>65 | 255.<br>20 | 59.<br>90 | 174.<br>35 | 54.4<br>9 | 249.<br>64 | 46.<br>32 | 244.<br>13 | 78.<br>07 | 209.<br>31 | 55.8<br>2 | 162.<br>68 | 41.<br>22 | 180.<br>62 | 42.<br>29 | 203.<br>78 | 61.<br>20 | 144.<br>14 | 33.3<br>5 | 177.<br>13 | 56.4<br>9 |
| Scotopic b-wave time | 54.7<br>3 | 8.9<br>3 | 53.6<br>7 | 6.3<br>0 | 48.5<br>7 | 7.3<br>9 | 52.8<br>7 | 9.74 | 56.7<br>1 | 5.3<br>9 | 46.3<br>3 | 3.5<br>9 | 59.1<br>4 | 9.58 | 53.2<br>7 | 6.8<br>0 | 47.0<br>0 | 3.7<br>1 | 52.7<br>5 | 5.8<br>0 | 46.6<br>6 | 5.33 | 50.3<br>8 | 6.62 |
| Photopic a- | 9.20 | 3.9 | 9.58 | 3.7 | 8.09 | 2.5 | 8.03 | 3.79 | 13.1 | 4.1 | 6.61 | 3.2 | 7.33 | 2.96 | 7.10 | 3.0 | 5.40 | 2.0 | 7.66 | 2.7 | 9.18 | 2.95 | 5.81 | 1.41 |

| wave<br>amplitude |  | 8 |  | 6 |  | 7 |  |  | 7 | 3 |  | 4 |  |  |  | 3 |  | 6 |  | 8 |  |  |  |  |
| --- | --- | --- | --- | --- | --- | --- | --- | --- | --- | --- | --- | --- | --- | --- | --- | --- | --- | --- | --- | --- | --- | --- | --- | --- |
| Photopic a-wave time | 18.0<br>4 | 3.1<br>5 | 17.9<br>0 | 3.4<br>8 | 15.2<br>1 | 1.1<br>7 | 18.1<br>5 | 2.99 | 17.6<br>6 | 3.5<br>7 | 16.2<br>9 | 2.0<br>4 | 17.1<br>6 | 2.51 | 9.96 | 2.8<br>7 | 18.3<br>9 | 3.5<br>6 | 17.3<br>1 | 1.8<br>4 | 17.7<br>5 | 2.30 | 15.9<br>3 | 1.46 |
| Photopic b-wave amplitude | 52.4<br>7 | 15.<br>49 | 79.2<br>6 | 20.<br>72 | 77.4<br>2 | 20.<br>52 | 52.0<br>5 | 13.5<br>8 | 86.9<br>5 | 19.<br>58 | 84.1<br>2 | 21.<br>00 | 59.2<br>2 | 12.1<br>3 | 39.9<br>7 | 11.<br>77 | 61.8<br>8 | 27.<br>77 | 60.4<br>3 | 15.<br>77 | 38.8<br>9 | 9.70 | 65.6<br>6 | 11.5<br>2 |
| Photopic b-wave time | 55.6<br>0 | 6.3<br>4 | 54.1<br>8 | 6.8<br>6 | 47.2<br>0 | 3.2<br>1 | 50.6<br>5 | 5.28 | 64.4<br>1 | 9.8<br>8 | 47.1<br>1 | 3.8<br>8 | 53.3<br>8 | 6.34 | 50.0<br>0 | 3.2<br>2 | 54.8<br>9 | 3.2<br>6 | 50.1<br>8 | 3.7<br>7 | 46.6<br>4 | 5.10 | 49.5<br>7 | 4.42 |
| PKCa | - | - | - | - | 490<br>5.67 | 107<br>1.4<br>8 | - | - | - | - | 324<br>9.65 | 105<br>7.7<br>1 | - | - | - | - | 324<br>3.30 | 697<br>.83 | - | - | - | - | 155<br>6.59 | 956.<br>06 |
| *Il1b* | - | - | - | - | 0.00<br>11 | 0.0<br>002<br>9 | - | - | - | - | 0.00<br>054 | 0.0<br>001<br>7 | - | - | - | - | 0.00<br>072 | 0.0<br>002<br>5 | - | - | - | - | 0.00<br>158 | 0.00<br>074 |
| *Nf-kb* | - | - | - | - | 0.00<br>171 | 0.0<br>007 | - | - | - | - | 0.00<br>122 | 0.0<br>002<br>9 | - | - | - | - | 0.00<br>118 | 0.0<br>002<br>4 | - | - | - | - | 0.00<br>105 | 0.00<br>026 |
| *Icam1* | - | - | - | - | 0.00<br>318 | 0.0<br>013<br>4 | - | - | - | - | 0.00<br>161 | 0.0<br>001<br>9 | - | - | - | - | 0.00<br>211 | 0.0<br>004<br>3 | - | - | - | - | 0.00<br>168 | 0.00<br>065 |
| *Vegfa* | - | - | - | - | 0.75<br>476 | 0.1<br>585<br>6 | - | - | - | - | 0.62<br>294 | 0.0<br>850<br>4 | - | - | - | - | 0.68<br>131 | 0.1<br>212<br>2 | - | - | - | - | 0.70<br>578 | 0.08<br>443 |

While this study provides compelling evidence of *APOE4*’s interaction with a 40% fat WD, future research could explore whether a 60% fat diet accelerates diabetic retinopathy in these mice. Longitudinal studies investigating the progression of diet-induced effects, along with interventions targeting specific inflammatory pathways, may offer more profound insights. Additionally, examining other cellular contributors to vascular dysfunction in *APOE4* mice could further elucidate the broader implications of this genotype on retinal health.

In conclusion, this study highlights the significant impact of dietary fat content on systemic metabolism, retinal function, and neuroinflammation in *APOE4* mice. While CD offers some metabolic benefits, it is not without its drawbacks, particularly in retinal health. These results highlight the importance of refining dietary strategies to balance systemic and retinal outcomes, especially in genetically vulnerable populations. These findings reinforce the necessity of genotype-specific nutritional strategies to mitigate metabolic and neurodegenerative risks associated with *APOE4*.

## Methods

### Study animals

Humanized *APOE*-knock-in mice were engineered by substituting the mouse Apoe gene’s exons 2, 3, and a significant portion of exon 4 with the human *APOE* gene sequence, including its exons 2, 3, and 4. The animals were homozygous for either *APOE3* (3/3) or *APOE4* (4/4) alleles and will be referred to as *APOE3* and *APOE4* mice, respectively. The mice were housed at the Eugene and Marilyn Glick Eye Institute, Indiana University, Indianapolis, under standard physiological conditions, including a 12-hour light/dark cycle with unrestricted access to food and water. The study received approval from the Institutional Animal Care and Use Committee (IACUC) and was conducted in accordance with the guidelines of the Association for Research in Vision and Ophthalmology and the National Institutes of Health regarding the ethical use of animals in research. Both male and female mice were included in all experiments.

### Dietary intervention

At approximately three months of age (13 weeks post-weaning), the mice were assigned to one of two diets: a high-fat Western diet (WD) comprising 40% kcal from fat, 43% kcal from carbohydrates, and 17% kcal from protein (4.7 kcal/g; D12079B, Research Diets, Inc., NJ, USA) or a low-fat control diet (CD) containing 10% kcal from fat, 73% kcal from carbohydrates, and 17% kcal from protein (3.9 kcal/g; D14042701, Research Diets, Inc., New Brunswick, NJ, USA). The diets were isocaloric. Experiments were conducted at three key time points: 21-26 weeks (∼2 months on diet), 39-43 weeks (∼6 months on diet), and 65-70 weeks (∼12 months on diet). Mice were euthanized at 65-70 weeks (15-16 months). Data from these three dietary intervention durations were compared between the WD and CD groups.

### Body weight and glucose tolerance assessment

Body weight was monitored monthly following the dietary intervention’s onset. Blood glucose levels were measured using an AlphaTRAK 2 glucometer (Zoetis, Parsippany, NJ, USA) with samples collected from the tail vein. An intraperitoneal glucose tolerance test (IPGTT) was performed after a 4–5-hour fasting period during daylight hours. Mice were administered glucose intraperitoneally (1g/kg body weight, Sigma-Aldrich, St. Louis, MO, USA), and blood glucose levels were recorded at 10, 20, 30, 60, 90, and 120 minutes post-injection.

### Plasma lipid profiling

After 12 months of dietary intervention, mice were euthanized under deep anesthesia, and whole blood was collected immediately following bilateral enucleation. Blood was obtained via the retro-orbital sinus using potassium EDTA-coated microvette capillary tubes (Sarstedt, Inc., Newton, NC, USA) to prevent coagulation. Blood samples were kept on ice and centrifuged at 2000 × g for 10 minutes at 4°C to separate plasma. The resulting plasma was aliquoted and stored at -80°C until analysis. Plasma lipid levels, including low-density lipoprotein (LDL), high- density lipoprotein (HDL), total cholesterol, triglycerides (TG), and free fatty acids (FFA), were measured using an automated clinical chemistry analyzer at the Indiana University School of Medicine’s Translational Research Core Facility.

### Optical coherence tomography (OCT) and fundus imaging

Retinal structural analysis was conducted using the Micron IV OCT system (Phoenix Technology Group, Bend, OR, USA). Mice were anesthetized via intraperitoneal injection of ketamine (88 mg/kg of body weight) and xylazine (12 mg/kg of body weight) and given topical 1% tropicamide/2.5% phenylephrine (ImprimisRX, Nashville, TN, USA) for pupil dilation. Eyes were lubricated using Gonak® or Hypromellose 2.5% solution (Akorn, Lake Forest, IL, USA). Once the pupils were dilated, the mice were placed on a stage of the Micron IV OCT system, and OCT-fundus images were captured, centering scans on the optic nerve. Both eyes were scanned, and retinal thickness was quantified using InSight software (phoenixmicron.com).

### Fluorescein angiography (FA)

For FA imaging, anesthetized mice were injected intraperitoneally with a 5 ml/kg dose of 2.5% fluorescein sodium solution. Retinal vasculature images were obtained under blue light illumination (490 nm) within 2 to 3 minutes of full vascular filling. FA was carried out using Micron IV OCT system, and images were analyzed for vein width, tortuosity, and vascular and avascular areas using MATLAB software (matlab.mathworks.com), employing quantification methods adapted from Mezu-Ndubuisi (2016).^59^

### Optomotor reflex (OMR) testing

Visual acuity and contrast sensitivity were assessed using the OptoMotry system (Cerebral Mechanics Inc., Lethbridge, Canada). Mice were placed in a centrally positioned arena surrounded by rotating grating stimuli of varying spatial frequencies. The setup was dimly lit so that the mouse maintains its focus on the moving stripes. Visual acuity was measured by presenting stripes with different spatial frequencies (the number of stripes per degree of visual angle), to identify the highest frequency the mouse could still follow. For contrast sensitivity, the contrast of the stripes was varied while keeping the spatial frequency constant. The direction of the visual stimulus was also changed (clockwise and anticlockwise) to observe mouse’s reflexive neck movements (from temporal to nasal), which reflected visual response of both eyes. Tracking responses were analyzed to determine visual acuity, measured in cycles per degree (cpd), and contrast sensitivity, expressed as the lowest detectable contrast level.

### Electroretinography (ERG)

ERG recordings were performed to evaluate retinal function under scotopic (dark-adapted) and photopic (light-adapted) conditions. Mice were dark-adapted overnight before anesthesia and pupil dilation (as mentioned above). After pupils were dilated, Gonak® or Hypromellose 2.5% solution was used to keep the eyes moist. A reference electrode was inserted sub-dermally between the eyes, while a ground electrode was inserted at the tail base. Gold loop electrodes were placed on both corneas. ERG signals were recorded using an LKC NGIT-100 (LKC Technologies, Inc., Gaithersburg, MD, USA) under different light intensities. Scotopic and photopic light responses were elicited using 2.5 candela-seconds per square meter (cd.s/m^2^) and 10 cd.s/m^2^ white light flashes, respectively; a 10-minute light adaptation preceded photopic exposure. a- and b-wave amplitudes, as well as implicit times, were analyzed using LKC EMWIN software (www.LKC.com).

### Quantitative real-time PCR (qRT-PCR)

Following 12 months of dietary intervention, mice were euthanized, and retinas were isolated post-enucleation. Retinal tissue was extracted using the Trizol-chloroform method (Thermo Fisher Scientific, Waltham, MA, USA), and cDNA was synthesized from 1μg RNA using the SuperScript Vilo Kit (Thermo Fisher Scientific). Gene expression was analyzed using qRT-PCR (Viia7, Thermo Fisher Scientific) with TaqMan Fast Universal Master Mix (Thermo Fisher Scientific) and gene-specific primers. *Il1b* (gene for Interleukin-1 beta, Mm00434228_m1), *Nf*κ*b* (gene for nuclear factor kappa-light-chain-enhancer of activated B cells, Mm00476361_m1), *Vegfa* (gene for vascular endothelial growth factor A, Mm00437306_m1)*, Icam-1* (gene for intercellular adhesion molecule 1, Mm00516023_m1) was quantified relative to the housekeeping gene *Tbp (*gene for TATA-box binding protein, Mm00446973_m1).

### Immunofluorescence staining

After 12 months on diet, eyes were fixed in 4% paraformaldehyde for 20 minutes at room temperature, washed in phosphate-buffered saline, and sectioned using a vibratome. Sections were incubated overnight at 4°C with primary antibodies against GFAP-1 (1:200, Cell Signaling Technology, Danvers, MA, USA) and PKCα (1:200, Sigma-Aldrich), followed by appropriate secondary antibody incubation. VECTASHIELD Antifade Mounting Medium with DAPI (Vector Laboratories, Burlingame, CA, USA) was used for nuclear staining. Images were captured using a Zeiss confocal microscope (LSM 700, Carl Zeiss Microimaging, Jena, Germany). Fluorescence intensity for PKCα was analyzed with Fiji ImageJ (fiji.sc), by subtracting secondary antibody control images from the PKCα staining images.

### Statistical analysis

Data analysis was conducted using GraphPad Prism 10.0.1 (GraphPad Software, CA, USA) and reported as Mean ± SEM. Statistical significance was set at p<0.05, with significance levels denoted as *p<0.05, **p<0.01, ***p<0.001, and ****p<0.0001. Data comparisons were performed using two-way ANOVA with Tukey’s post hoc test, while PKCα, *Il1b, Nf*κ*b, Vegfa,* and *Icam-1* quantifications were analyzed via one-way ANOVA with Tukey’s post hoc analysis.

## Abbreviations

AD: Alzheimer’s disease
LOAD: Late-onset Alzheimer’s disease
APOE3: Apolipoprotein
E3 APOE4: Apolipoprotein E4 T2D: Type 2 diabetes
HFDs: High-fat diets
WD: Western diet CD: Control diet
IPGTT: Intraperitoneal glucose tolerance test OCT: Optical coherence tomography
FA: Fluorescein angiography
ERG: Electroretinogram OMR: Optomotor reflex

### Acknowledgments

We thank Dr. Evan Cornett, Dr. Jeffrey Elmendorf, and Dr. Amelia Linnemann from Indiana University School of Medicine for their valuable suggestions and guidance.

## Authors’ contributions

SDA: Writing–original draft, writing–review & editing, conceptualization, software, validation, formal analysis, and investigation. QL: Writing–review & editing, validation, software, formal analysis, and investigation. GH: Software, validation, formal analysis, investigation, and writing–review & editing. NM: Investigation, writing, review & editing. TWC: Supervision, resources, and writing–review & editing. ALO and BTL: Resources and writing–review & editing. AB: Conceptualization, resources, writing–review & editing, supervision, project administration, and funding acquisition.

## Data availability

The data will be available from the corresponding author upon request.

---

Supplemental Figure 1: Increased plasma lipids in mice fed a WD.

(A) Low-density lipoprotein cholesterol (LDL) levels were significantly increased in *APOE3* mice fed a WD compared to CD, while *APOE4* mice showed a non-significant upward trend. (B) Triglyceride (TG) levels were elevated in both *APOE3* and *APOE4* WD-fed groups, though differences did not reach statistical significance. (C) Total cholesterol levels were significantly higher in *APOE3* WD-fed mice compared to *APOE3* CD controls, with modest increases in *APOE4* mice. (D) Free fatty acid (FFA) levels displayed a complex pattern: *APOE3* WD mice exhibited a non-significant reduction in FFA compared to *APOE3* CD, while FFA levels were significantly elevated in *APOE3* CD and *APOE4* WD mice compared to *APOE4* CD controls. (E) High- density lipoprotein cholesterol (HDL) levels were significantly increased in *APOE3* WD-fed mice compared to *APOE3* CD, with non-significant increases observed in *APOE4* mice. Values are expressed as mean ± S.E.M. (n= 5 mice/ group). One-way ANOVA with Tukey’s test. *p<0.05, **p<0.01, ***p<0.001.

