## Supplementary figures and images for "Long-term Dietary Fat Intervention Affects Retinal Health in APOE Mice"

### Supplemental Figure 1

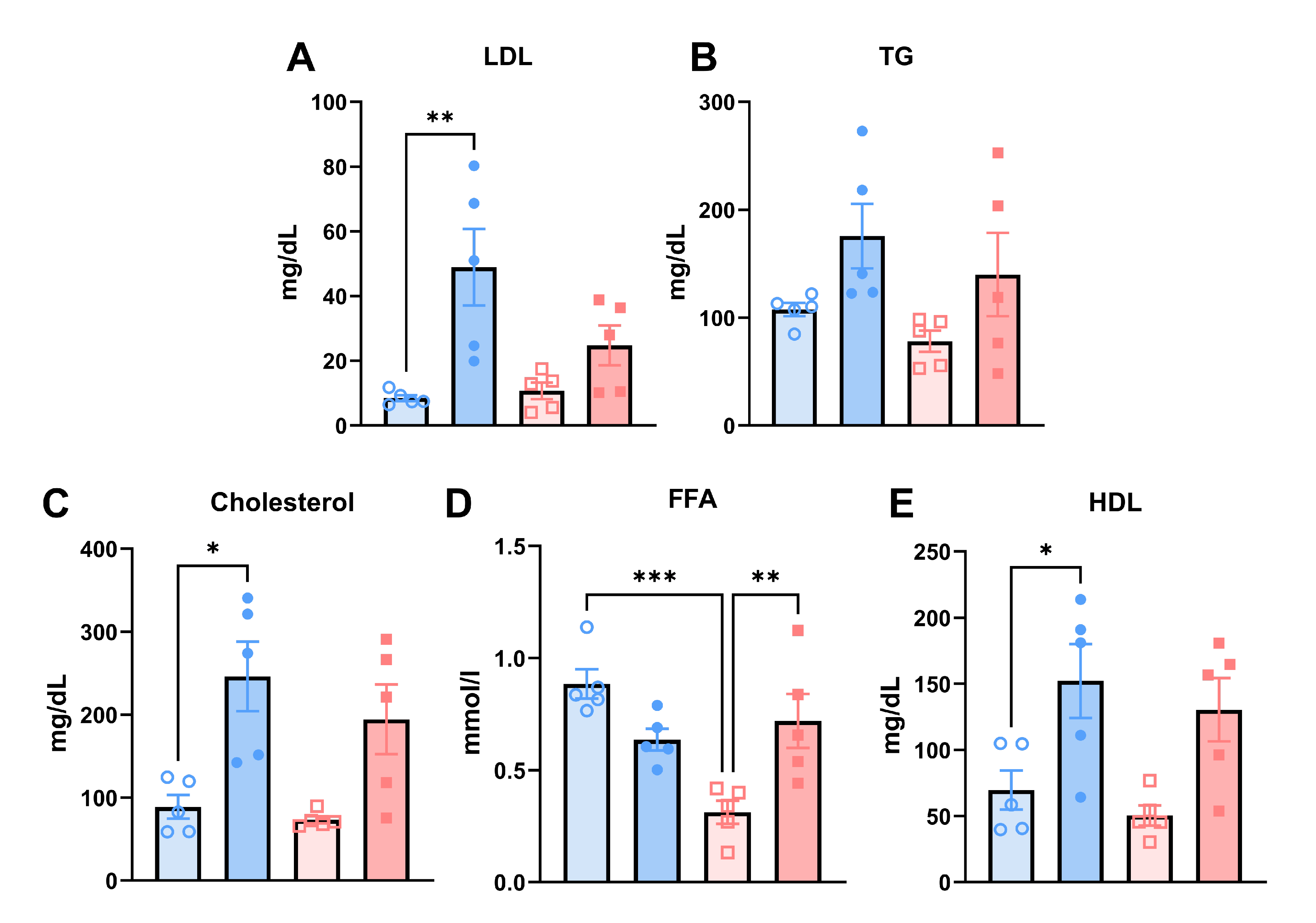
